# A high-throughput 3D bioprinted cancer cell migration and invasion model with versatile and broad biological applicability

**DOI:** 10.1101/2021.12.28.474387

**Authors:** MoonSun Jung, Joanna N. Skhinas, Eric Y. Du, M.A. Kristine Tolentino, Robert H. Utama, Martin Engel, Alexander Volkerling, Andrew Sexton, Aidan P. O’Mahony, Julio C. C. Ribeiro, J. Justin Gooding, Maria Kavallaris

**Affiliations:** Children’s Cancer Institute, Lowy Cancer Research Center, UNSW Sydney, NSW, 2031, Australia; Australian Center for NanoMedicine, UNSW Sydney, NSW 2031, Australia; School of Women and Children’s Health, Faculty of Medicine and Health, UNSW Sydney, NSW, 2031, Australia; School of Chemistry, UNSW Sydney, NSW 2052, Australia; Inventia Life Science Pty Ltd, Sydney, NSW 2015, Australia

**Keywords:** 3D bioprinting, high-throughput platform, *in vitro* 3D model, tunable hydrogels, cell migration, metastasis

## Abstract

Understanding the underlying mechanisms of migration and metastasis is a key focus of cancer research. There is an urgent need to develop *in vitro* 3D tumor models that can mimic physiological cell-cell and cell-extracellular matrix interactions, with high reproducibility and that are suitable for high throughput (HTP) drug screening. Here, we developed a HTP 3D bioprinted migration model using a bespoke drop-on-demand bioprinting platform. This HTP platform coupled with tunable hydrogel systems enables (i) the rapid encapsulation of cancer cells within *in vivo* tumor mimicking matrices, (ii) *in situ* and real-time measurement of cell movement, (iii) detailed molecular analysis for the study of mechanisms underlying cell migration and invasion, and (iv) the identification of novel therapeutic options. This work demonstrates that this HTP 3D bioprinted cell migration platform has broad applications across quantitative cell and cancer biology as well as drug screening.

## Introduction

Metastasis is the leading cause of cancer-related deaths and thus an understanding of the underlying molecular and cellular processes continues to be a key focus of cancer research. Tumor dissemination is a multistep process. It involves cancer cell migration and invasion through the extracellular matrix (ECM) to allow entry into adjacent tissues, blood, and lymphatic vessels (Novikov et al., 2021). The migration and invasion of cancer cells is a dynamic activity of cells, regulated by integrins, matrix-degrading enzymes, cell–cell adhesion molecules and cell–cell communication in living tissues (Friedl and Wolf, 2003). However, current understanding of the molecular mechanisms underlying cell migration and invasion remains heavily dependent on cancer cells grown on flat 2-dimensional (2D) plastic. These 2D model systems are frequently used in high throughput (HTP) drug discovery although they do not always accurately predict drug response. A scarcity of appropriate screening platforms that can be directly translatable from *in vitro* to *in vivo* is a major contributor that hinders the development of drugs specifically targeting cancer metastasis. As such, there is a growing consensus regarding the use of 3D cell models to better mimic physiological cell-cell and cell-ECM interactions as more suitable approaches for identifying novel inhibitors disrupting cell migration than 2D cell culture.

The classic *in vitro* 3D cell migration and invasion models are the Boyden chamber transwell-based assays. In the transwell assays, cells migrate through a physical barrier, including membrane pores and ECM-mimicking materials, toward a chemo-attractant gradient. The simplicity and low-cost for set-up make these methods an excellent tool in cancer research. The transwell assays is often low throughput and gives only an endpoint readout. More automated fabrication of 3D cell models with higher throughput and real-time measurement of cell movement would be a significant advance over the transwell-based approaches. Additional advances could come from improvements in the tunability of the ECM mimic as this is an important component of any 3D cell migration and invasion assays, including the transwell assays. Most 3D cell model systems currently use naturally derived hydrogels, such as collagen and Matrigel. Of those, Matrigel, a basement membrane matrix composed of a complex mixture of various proteins has been widely used with a variety of cell types. It is recognized as a “gold standard” scaffold for numerous *in vitro* 3D cell culture applications. Despite Matrigel being a powerful resource as an ECM mimic for the study of cell biology for decades, batches-to-batch variability and challenges in tuning the chemical and physical properties of Matrigel lead to a lack of reproducibility and hampers unambiguous determination of interactions between cell, physical, and biochemical ECM factors, including stiffness and integrin-binding proteins (Kratochvil et al., 2019, Fang and Eglen, 2017). Thus, it is important to develop matrix scaffolds that are more reproducible and tunable for their mechanical, chemical, physical, and biological properties for 3D cell models. Accordingly, an automated 3D model fabrication combined with tunable matrix materials would increase the reliability of *in vitro* cell migration and invasion experiments.

Numerous biomaterial-based fabrication technologies, including 3D bioprinting, have recently been developed for the creation of 3D constructs using ECM mimics that have more tunable properties than Matrigel. 3D bioprinting provides a new approach to fabricate cell-laden matrices in a well-controlled manner over the positioning of cells and synthetic hydrogels (Moroni et al., 2018). Thus, it has emerged as a promising tool to engineer complex 3D tumor-microenvironments *in vitro* to study various biological processes including cell migration and invasion. Several 3D bioprinted metastasis models developed using tunable hydrogel systems have recently been reported (Ding et al., 2019, Meng et al., 2019, Zhou et al., 2016). Though, only a few studies have demonstrated a 3D bioprinted platform with HTP capability and versatility for multiple biological applications.

Recently we developed an enabling technology consisting of a bespoke drop-on-demand 3D bioprinter, which employs a fly-by printing logic that allows for the ejection of ink droplets at a constant and rapid speed in multi-well plates (Utama et al., 2021, Utama et al., 2020). In our recent work, we demonstrated that the hydrogel system of a 4-arm poly(ethylene glycol) maleimide (PEG-4MAL) bioink and a bis-thiol activator that instantly cross-links the polymer can be used as the 3D ECM-like hydrogels, due to the highly tunable mechanical and biofunctional properties (Lutolf et al., 2003), as well as their printability (Utama et al., 2021). In the present study, we leveraged the 3D bioprinting technology and tunable hydrogel system to develop a versatile HTP 3D bioprinting platform for studying cell migration and invasion. Herein as cell model systems, we selected a combination of cancer cell lines with non-invasive and invasive phenotypes, which are known to express distinct epithelial and mesenchymal markers (Liu et al., 2015, Havel et al., 2015, Swaminathan et al., 2011). We demonstrated that the platform can be utilized to identify cell-type specific ECM-like hydrogels by tuning the matrix stiffness and biochemical molecules, and capabilities for downstream phenotypic and molecular analysis *in situ* using non-invasive and invasive cancer cell models. Moreover, cell movement within the 3D constructs can be monitored, tracked and measured in real-time using the platform. The proposed bioprinted platform could serve as a powerful tool to understand the mechanisms of cancer cell migration and invasion and be used in HTP drug testing to discover drugs effective against cancer metastasis.

## Results

### Design of HTP bioprinting platform with tunable hydrogels

The 3D cell-laden hydrogel constructs were printed in multi-well plates via a two-droplet bioprinting process in which the first droplets containing PEG-4MAL bioink were printed, followed by printing of the second droplets of the bis-thiol activator mixed with cancer cells onto the bioink droplet for instant hydrogel formation as previously described (Utama et al., 2021). The process facilitated the generation of simple and highly reproducible 3D cancer models to study cell movement in well-defined ECM environments in a HTP manner. We demonstrated using the MCF7 breast cancer cell line that the PEG-4MAL hydrogel could successfully encapsulate cells as part of the bioprinting process and provide a platform for a stable 3D culture (Figure 1A). To assess the accuracy of the printing platform, we repeatedly printed MCF7 cells in 96 well plates and measured the cell growth for up to 7 days using the Alamar Blue metabolic assay. For each time point, we had 10 replicates per plate and similar absorbance values for each replicate were obtained, confirming the high repeatability in one print run (Figure 1B). Three independent print runs of the MCF7 cells were performed. We found that the absorbance readings of cell metabolic activity for technical replicates were very similar (Figure 1C), indicating that the HTP bioprinting platform can print and encapsulate cells in the 3D hydrogels with high reproducibility. Furthermore, by taking advantage of the HTP 3D bioprinting technology, the platform can generate up to 3 different tumor models from 3 different cell types (MDA-MB-231, MCF7 and H1299 cells) in the same multi-well plate with many replicates, enabling the testing of several cell types and hydrogels concurrently (Supplementary Figure 1).

**Figure 1.**
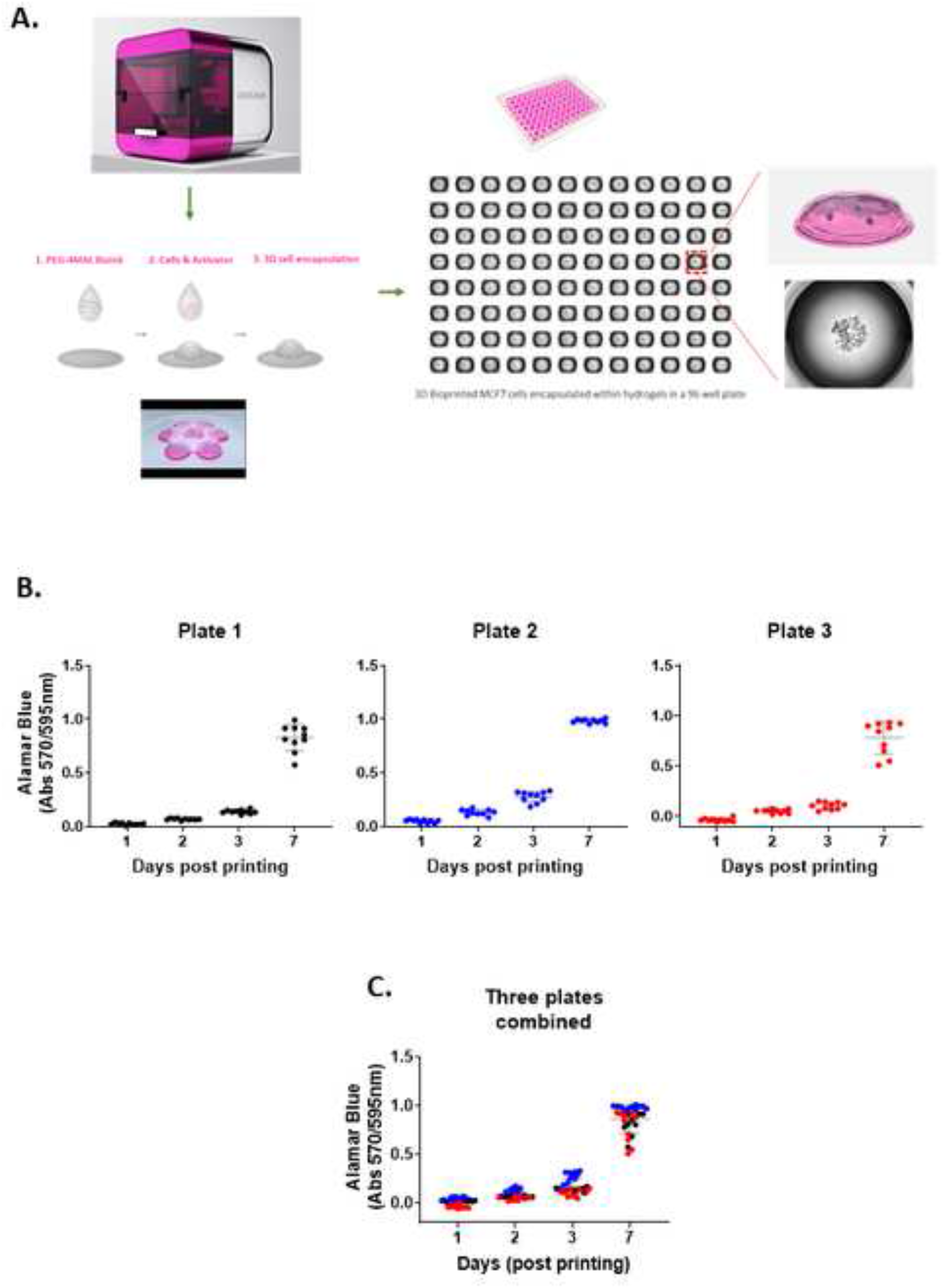
A 3D HTP bioprinting platform using tunable hydrogel systems. (A). A schematic of the bioprinting process. 3D cell models are generated using the bespoke drop-on-demand 3D bioprinter. Cancer cells are encapsulated within the hydrogel via a two-droplet system where first droplet contains a PEG-4MAL bioink and second droplet holds cells and MMP-sensitive activator. When the two droplets meet, the instant gelation happens, which enabling a rapid production of 3D cell models in a multi-well plate. (B). Repeatability and (C). Reproducibility of the 3D bioprinting platform. MCF7 cells were bioprinted in a 96 well plate and cell proliferation rate was measured at day 1, 3, 5 and 7 by Alamar Blue assay. 10 replicate wells were used for each time point per plate. Three independent runs were performed.

### Biocompatibility of 3D bioprinted hydrogels

Next, to investigate the biocompatibility of the bioprinted hydrogels, we selected 4 different PEG-4MAL hydrogel combinations, which exhibited two different levels of stiffness, 0.7 kPa and 1.1 kPa. Each PEG-4MAL bioink was decorated with or without RGD (arginine-glycine-aspartic acid) cell adhesion peptides and crosslinked with MMP (matrix metalloproteinase)-cleavable activator; 0.7 kPa, 0.7 kPa+RGD, 1.1 kPa, and 1,1 kPa+RGD hydrogels. These hydrogel combinations were chosen to demonstrate the effect of mechanical (stiffness) and biological molecules (the cell adhesion peptides) of a tumor-like microenvironment on cell responses (growth and migration). We chose four epithelial cancer cell lines that are known to have differing migratory and invasive properties, including two variants of breast cancer cell lines, MCF7 (basal-like; non-invasive) and MDA-MB-231 (triple-negative; invasive), and two other invasive cell lines, HEY (high-grade serous ovarian cancer) and H1299 (lung cancer).

To determine cell viability, each cancer cell line was bioprinted and encapsulated within the hydrogel systems in 96 well plates and cultured for 7 days post-printing. All four cell lines remained highly viable in those four hydrogel conditions (Figure 2A). Our data shows that the growth of cancer cells was dependent on various parameters, such as cell type, stiffness and the presence or the absence of the RGD peptides. While MCF7 non-invasive breast cancer cells appeared to grow similarly in all 4 hydrogel conditions, MDA-MB-231, an invasive variant of breast cancer, was shown to be highly proliferative only in the presence of the cell adhesion peptides, RGD, irrespective of the different stiffness of the hydrogels (Figure 2B). Similar to MDA-MB-231, H1299, an invasive lung cancer cell displayed high metabolic activity in the hydrogels incorporated with RGD, 0.7 kPa+RGD or 1.1 kPa+RGD hydrogels. In contrast, the growth of HEY cells, derived from aggressive high-grade serous ovarian cancer, was observed in both 0.7 kPa hydrogels with or without RGD, while in the 1.1 kPa hydrogels, the presence of RGD seemed to be vital for their growth (Figure 2B). Altogether, these data suggest that while bioprinted PEG-4MAL hydrogel systems are highly biocompatible, and each cancer cell type requires distinct matrix components for their growth.

**Figure 2.**
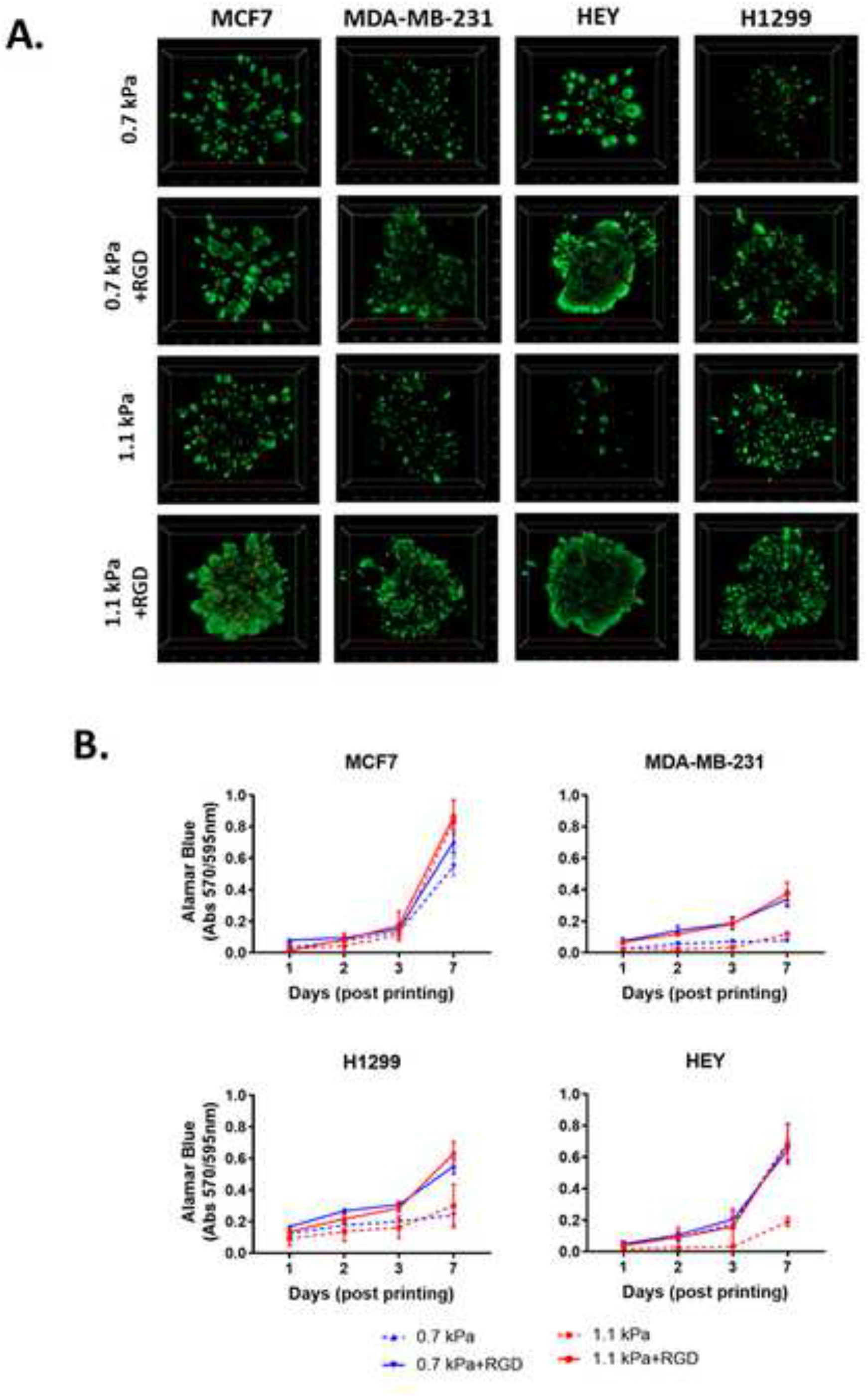
Biocompatibility of 3D bioprinted hydrogels. (A). Viability of 3D bioprinted cancer cells. Cancer cells were bioprinted with each hydrogel combination (0.7 kPa±RGD or 1.1 kPa±RGD) in 96 well plates and cultured for 7 days. Cells were stained with calcein-AM (green; live)/ethidium homodimer (red; dead) Live/Dead Assay and z-stack 3D images were taken at day 7 post-printing (5X objective). All experiments were repeated three times. (B). Cell proliferation of 3D bioprinted cancer cells. Each cancer cell line was bioprinted and encapsulated in one of the four hydrogel combinations and was cultured for up to 7 days. Growth rate was measured using Alamar Blue assay at day 1, 2, 3 and 7 post-printing. All experiments were repeated three times.

### Morphology of cancer cells encapsulated in ECM-like hydrogels

3D cell morphology can be used to define cellular behavior and function and predict the malignant potential of cells (Benton et al., 2011, Weigelt and Bissell, 2008). Thus, we next determined the impact of the matrix conditions on cell morphology by modulating hydrogel stiffness and/or adhesion peptides. Cells were bioprinted and encapsulated within the hydrogels, which were optimal for their growth as determined in Figure 2. In parallel, cells were encapsulated within Matrigel, a widely used biomaterial, as a reference control of their 3D morphology. As expected, the two variants of breast cancer cells showed distinct morphology when cultured within Matrigel. MCF7 cells were found to form multiple spheroids in Matrigel, whereas the MDA-MB-231 cells invaded through the Matrigel and exhibited stellate and protrusion morphology at day 7 (Figure 3A, Movies 1 and 2), in line with previous observations (Benton et al., 2011). When MCF7 cells were bioprinted in any of the four hydrogels, cells formed spherical structures and were predominantly proliferative from a single colony rather than migratory (Figure 3B, Movie 3). Interestingly, we found that the spheroid morphology appeared to vary between the hydrogel conditions, in terms of their size and roundness, a measure of how close the shape of the 2D spheroid image approaches a circle (Amaral et al., 2017). In the presence of ECM mimics that have conjugated RGD peptides (0.7 kPa+RGD and 1.1 kPa+RGD hydrogels), the generated MCF7 3D models displayed spherical structures that were larger in size, irregular, and had less-round shapes (Figure 3B). In contrast, the mean size and roundness of 3D bioprinted MCF7 models in the absence of RGD peptides were similar to those obtained with Matrigel (size: 0.18±0.10 μm^2^; roundness 0.86±0.089), especially the spheroids generated with 0.7 kPa hydrogel (size: 0.23±0.18 μm^2^; roundness: 0.80 ± 0.12) (Figure 3C). This suggests that despite the similar proliferation rate of MCF7 cells in all hydrogel conditions, the mechanical and biological characteristics of the matrix can affect cell morphology. MDA-MB-231 cells bioprinted in 0.7 kPa+RGD or 1.1 kPa+RGD hydrogels also showed the protrusion and network forming morphology and appeared to migrate through the hydrogels in a similar way to that observed in Matrigel (Figure 3 A and B, Movies 2 and 4). Moreover, the morphology seemed not to be affected by the hydrogel stiffness. In addition to the metastatic breast cancer cells, a similar morphologic pattern was observed in metastatic lung cancer (H1299) and ovarian cancer (HEY) cell lines (Figure 3B). The bioprinted 3D models for both H1299 and HEY cells had similar morphology in 0.7 kPa+RGD and 1.1 kPa+RGD hydrogels as they did in Matrigel. Yet, when HEY cells were bioprinted in 0.7 kPa hydrogels without RGD peptide, the cells displayed spheroid-like morphology (Figure 3D). Hence, these indicate the matrix components, such as cell adhesion molecules can alter morphology of cells grown on the hydrogels.

**Figure 3.**
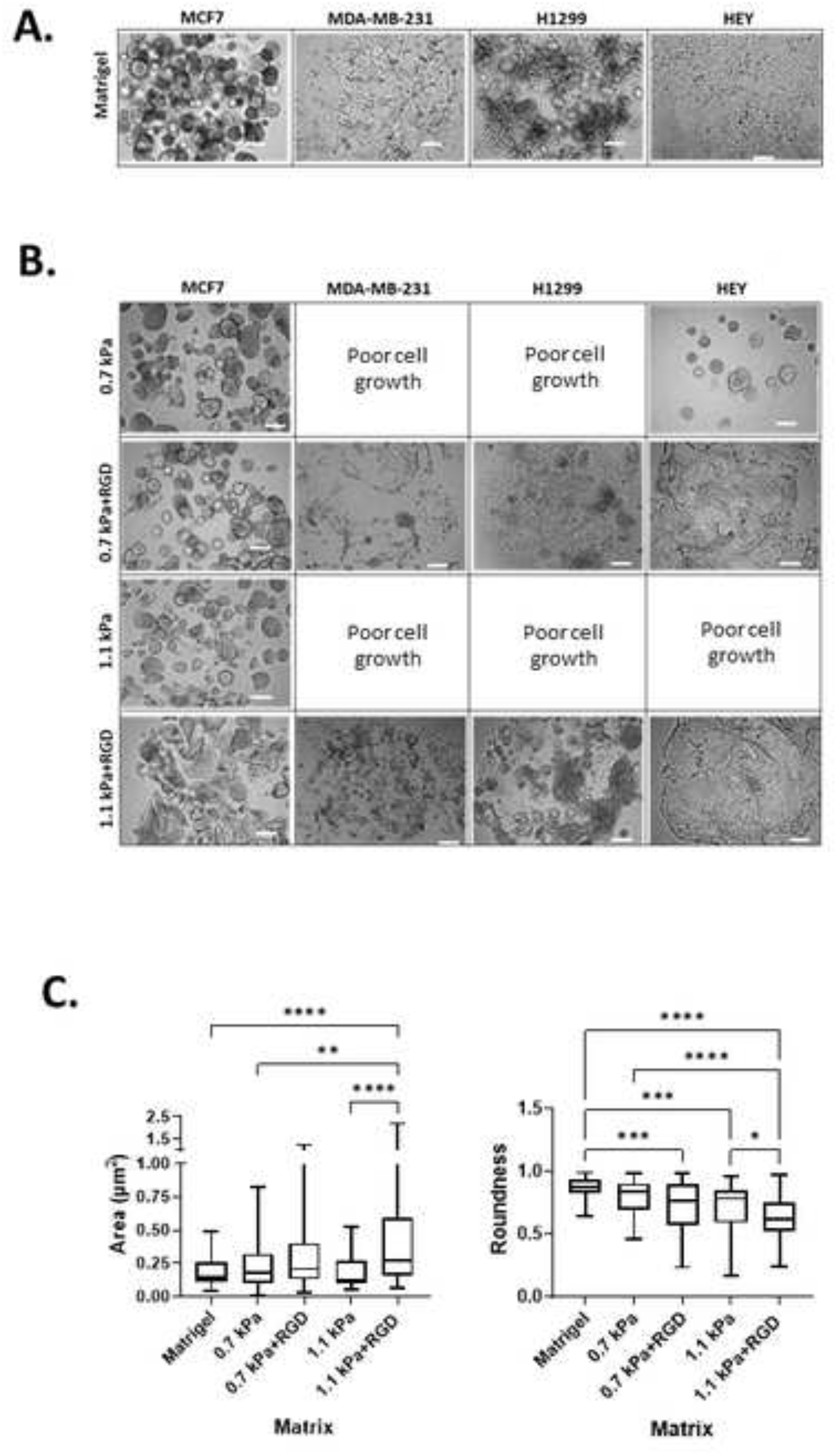
Cancer cell morphology in 3D bioprinted hydrogels. (A). 3D cell morphology in Matrigel. Each cell line was manually encapsulated in 2 μl of Matrigel in a 96 well plate. Cells were cultured for 7 days prior to taking bright-field imaging at a single plane (Scale bar: 200μm) (B). 3D cell morphology in bioprinted hydrogels. Each cell line was bioprinted with the hydrogel conditions optimal for their growth in a 96 well plate. Cells were cultured for 7 days prior to taking bright-field imaging at a single plane (Scale bar: 200μm). All experiments were repeated at least twice. (C). Analysis of MCF7 cell spheroids in Matrigel and bioprinted hydrogels. The size and roundness of 3D spheroids of MCF7 cells (at least 40 spheroids per condition; images in (A) and (B)) were analyzed and quantified using ImageJ Fiji. \**P*<0.05, \*\**P*<0.01, \*\*\**P*<0.001, \*\*\*\**P*<0.0001, One-way ANOVA with a post-hoc Tukey test for comparison between means.

### Expression of metastasis relevant genes and proteins in 3D bioprinted cancer cells

We next demonstrated the suitability of this platform to be used for the analysis of phenotypic markers relevant to cell migration and invasion in 3D cancer models. Epithelial-to-mesenchymal transition (EMT) has been implicated in cancer invasion and progression (Roche, 2018, Brabletz et al., 2018, Havel et al., 2015), which is associated with a loss of epithelial markers such as E-cadherin and a gain of mesenchymal markers such as vimentin. Bioprinted 3D models of each cell line in their optimized hydrogel conditions were cultured for 7 days, and then subjected to *in situ* immunofluorescent staining for the simultaneous detection of several proteins involved in cell migration including EMT process (E-cadherin and vimentin) as well as ECM-remodeling (MMP2 and MMP9). We confirmed that E-cadherin was predominantly expressed in MCF7 cells, while positive vimentin expression was detected in the invasive cancer cell lines, MDA-MB-231, H1299 and HEY, showing that the EMT phenotypic markers were retained in the 3D bioprinted cell models (Figure 4A). Interestingly, while positive expression of MMP2 and MMP9 was found in all 3D bioprinted models, the expression of both MMP2 and MMP9 appeared to be prominent in the invasive cell lines, MDA-MB-231, H1299 and HEY (Figure 4A).

**Figure 4.**
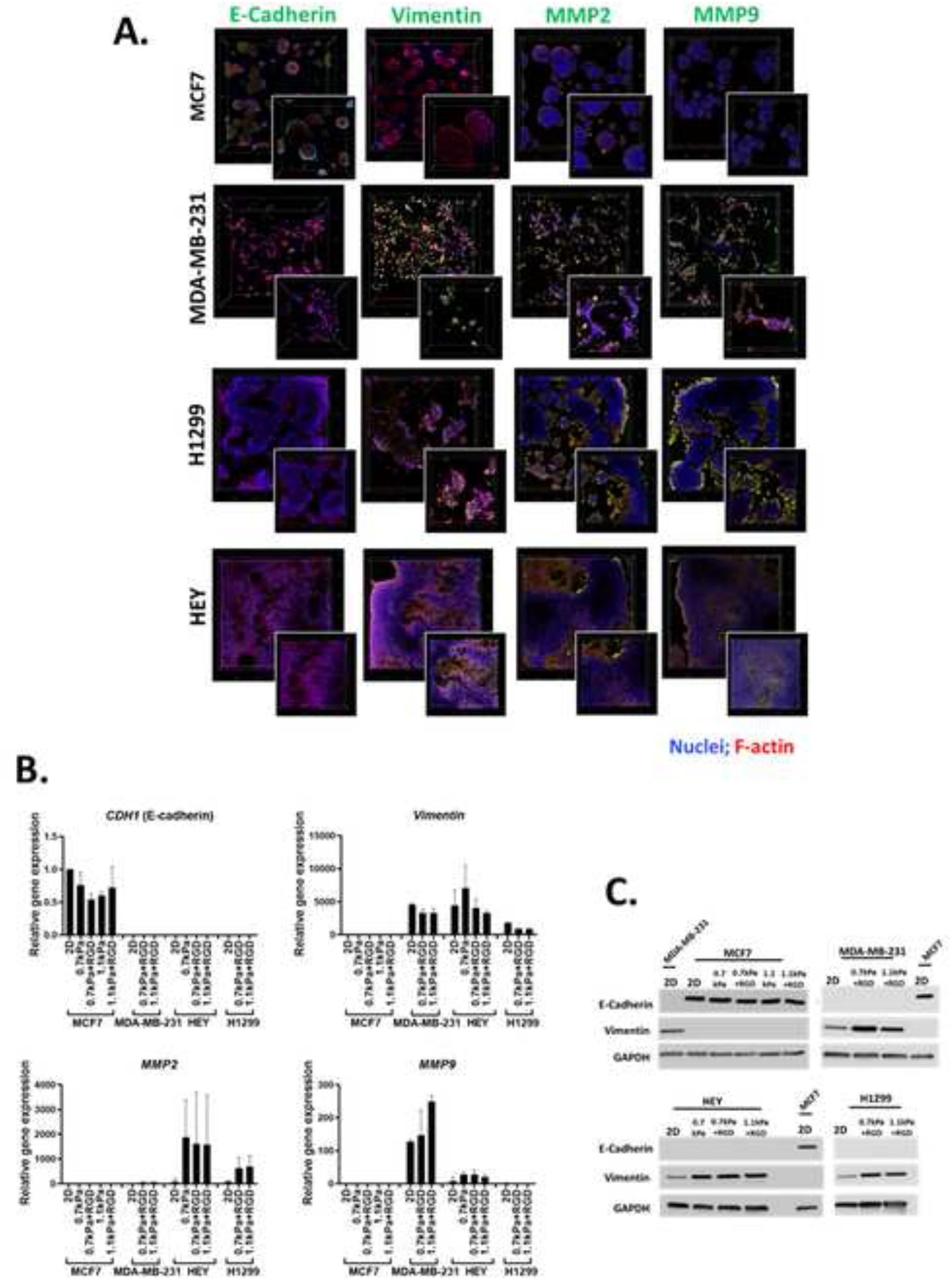
*In situ* microscopic and molecular analysis of phenotypic markers using the 3D bioprinting platform. (A). *In-situ* immunofluorescent images of migration related proteins. Each cell line was bioprinted in its optimum hydrogels and cultured for 7 days prior to fixation with 4 % paraformaldehyde. Cells were stained for F-actin (red), nuclei (blue) and migration/invasion relevant proteins (E-cadherin, vimentin, MMP2 and MMP9; green). Confocal microscopic images were taken (10X objective; Inset: Zoomed-in images). (B). Gene expression and (C) protein expression analysis using cells recovered from the 3D hydrogels via an enzymatic degradation approach. Retrieved cells were subjected to qPCR and Western blot for the migration/invasion relevant mRNA and protein expression levels respectively. All experiments were repeated at least twice.

Having shown the qualitative expression of migration and invasion associated proteins in the hydrogel embedded cells, we next sought to investigate the quantitative expression of the relevant genes and proteins. Due to the presence of MMP-sensitive peptides within the PEG-4MAL hydrogel system, cells can be readily retrieved from the hydrogels *via* proteolytic degradation. Using this hydrogel feature, we next determined the versatility of the bioprinting platform for detailed molecular and protein analysis. Cells bioprinted in 96 well plates were recovered from the hydrogels through enzymatic approaches after 7 days in culture. Expression levels of metastasis relevant genes and proteins in the cells recovered from hydrogels was determined using qPCR and Western blotting, respectively. MCF7 cells recovered from all 4 hydrogel conditions exhibited similar levels of E-cadherin *(CDH1* gene) gene and protein expression to their 2D culture counterpart (Figure 4B and C). As expected, vimentin was not expressed at the gene or protein levels in the MCF7 cells (Figure 4B and C). On the contrary, all the invasive breast, lung and ovarian cancer cells (MDA-MB-231, H1299 and HEY respectively) showed expression of vimentin but no E-cadherin expression in any hydrogel conditions (Figure 4B and C). Additionally, consistent with the *in situ* immunofluorescence image analysis (Figure 4A), gene expression of the ECM-remodeling proteases was prominently detected in the invasive cell lines, MDA-MB-231 and HEY cells, and H1299 while neither MMP2 nor MMP9 mRNA expression was detectable in MCF7 cells (Figure 4B). Despite the positive protein staining of MMP2 and MMP9 (Figure 4A), only MMP2 mRNA expression was detected in H1299 cells (Figure 4B), suggesting that mRNA levels are not always proportional to protein levels (Mehra et al., 2003). Altogether, these show that this 3D bioprinting platform employing PEG-4MAL hydrogel system can be a simple and versatile approach to be routinely used for molecular and *in situ* analysis to study cancer cell migration and invasion.

### 3D bioprinted platform for cell movement tracking in real-time

Although several methods have been developed for visualizing and analyzing cell migration in a 2D setting, no simple and straightforward cell tracking approach to quantify 3D cell movement has been established. Having demonstrated that invasive cancer cells retained their mesenchymal characteristics in the bioprinted hydrogels, we next sought to validate the potential of our bioprinting platform to be used as a methodology for studying cancer cell migration and invasion. To measure the dynamic movement of single cells within the bioprinted hydrogels, nuclei-labelled live cells were monitored for a period of 24 h on day 1 of post-printing using an automated widefield microscopic imaging system and then analyzed using object tracking. We quantified migratory properties including track length, displacement and mean speed of cell movement (Figure 5A). When comparing cell movement between the two breast cancer cell variants, MDA-MB-231 exhibited greater migratory behavior with a longer track length (P<0.0001) and faster movement (P<0.05) than MCF7 cells within the same hydrogel condition, 0.7 kPa+RGD (Figure 5B). In MDA-MB-231 cells, the hydrogel stiffness impacted migration behavior. Greater migratory behavior of MDA-MB-231 cells was observed within the stiffer hydrogels (1.1 kPa+RGD) compared to the softer hydrogel system, 0.7 kPa+RGD (Figure 5C). These findings demonstrate that the mechanical property of the hydrogels can affect cell migration behaviors.

**Figure 5.**
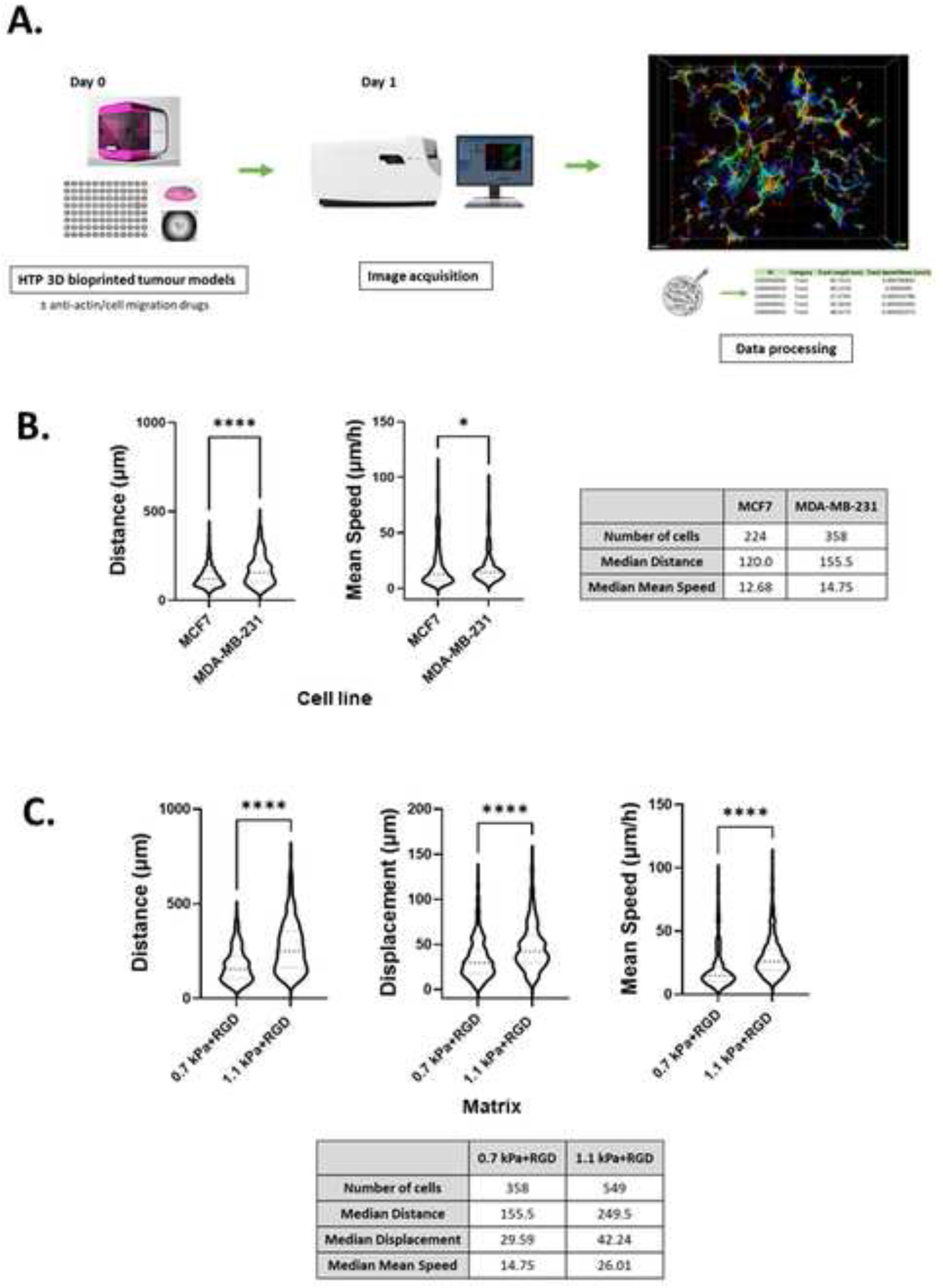
Tracking migratory behaviors of cancer cells in real-time using the HTP 3D bioprinting platform. (A). A schematic of the workflow of 3D cell movement tracking. 3D cell models were generated in a multi-well plate using the 3D bioprinting platform (Day 0) and cultured for 24 h with/without chemical inhibitors (Day 1). Cell movement within the 3D hydrogels was monitored and tracked for a period of 24 h from day 1 of post-printing. Tracks were normalized with respect to dead cell control wells. (B). Quantitation of migratory behavior of MCF7 and MDA-MB-231 cells within 0.7 kPa+RGD hydrogels. \**P*<0.05, \*\*\*\**P*<0.0001, Mann-Whitney test was performed. (C). Quantitation of migratory behavior of MDA-MB-231 cells within either 0.7 kPa+RGD or 1.1 kPa+RGD hydrogels. All experiments were repeated at least twice. \*\*\*\**P*<0.0001, Mann-Whitney test was performed.

### 3D bioprinted cells as a screening platform for pharmacological inhibitors of migration and invasion

We further determined the feasibility of our 3D bioprinting platform as a preclinical model for anti-metastatic drug testing. Here, we tested the effect of two known pharmacological inhibitors of cell migration, Y-27632 (a ROCK inhibitor) and blebbistatin (a global myosin inhibitor) on cell migratory behaviors within one of the hydrogels, 1.1 kPa+RGD. Once the 3D tumor models were bioprinted in 96 well plates, cells in each well were treated with either of the drugs for 48 h and tracked from 24 h to 48 h after printing. Using MDA-MB-231 cells, we successfully monitored 3D movement of cells treated with different drugs simultaneously and quantitate their track distance, displacement and mean speed of cell movement. As shown in Figure 6A, 3D migration of MDA-MB-231 cells was significantly impeded by the ROCK inhibitor (track distance P<0.001; displacement P<0.0001; mean speed P<0.0001) or blebbistatin (P<0.0001 for all parameters) compared to no drug-treated cells in 1.1 kPa+RGD hydrogel. The significant inhibitory effects on MDA-MB-231 cell movement were also observed in the softer gels (0.7 kPa+RGD; Supplementary Figure 2). For further validation, another invasive cancer cell line, H1299 bioprinted with 0.7 kPa+RGD hydrogels, was treated with either of the inhibitors. As observed in MDA-MB-231 cells, suppression of H1299 cell movement upon drug treatment could be monitored and quantitated using the platform (Figure 6B). Overall, the 3D bioprinted platform we propose here cannot only accurately measure cell migratory behaviors within the 3D matrices but also has application as a HTP preclinical anti-metastasis drug testing platform.

**Figure 6.**
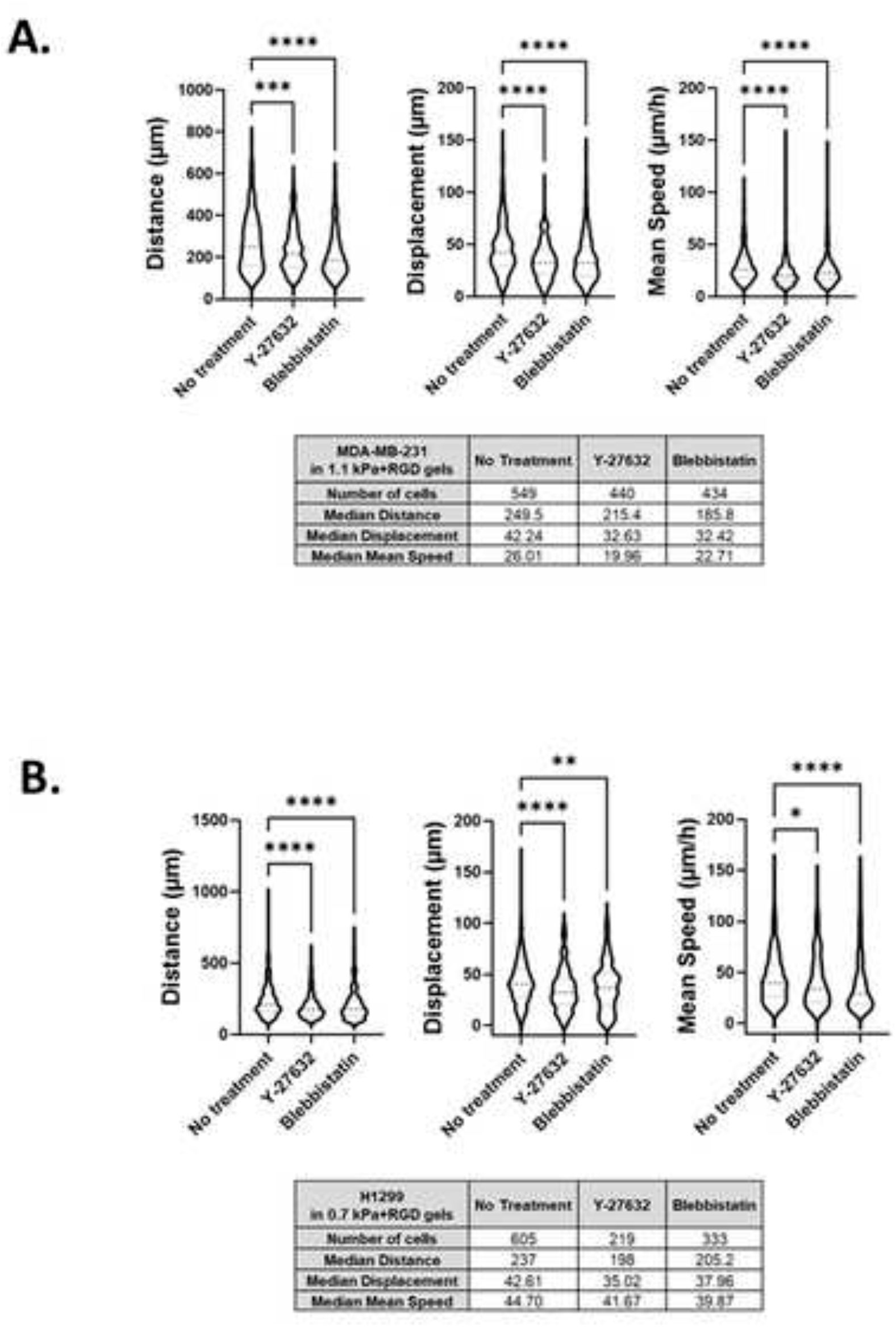
Screening of chemical inhibitors of cell movement using the 3D bioprinted platform. (A) and (B). Y-27632, a ROCK inhibitor and blebbistatin, a global myosin inhibitor, were treated in bioprinted cancer cells in 96 well plates. 3D cell movement of MDA-MB-231 (A) and H1299 (B) cells in the absence or presence of the inhibitors was compared and quantitated. All experiments were repeated at least twice. Kruskal–Wallis one-way analysis with a post hoc Dunn test was performed. \**P*<0.05, \*\**P*<0.01, \*\*\**P*<0.001, \*\*\*\**P*<0.0001

## Discussion

Despite recent advances in 3D bioprinting and numerous reports on its potential applications in cancer research (Hwang et al., 2021, Swaminathan et al., 2019, Meng et al., 2019, Heinrich et al., 2019, Zhou et al., 2016), there are few bioprinting platforms that are simple to use, capable of HTP and suitable for multiple biological assays with high cell viability. In this study, we have developed unprecedented capabilities for a 3D bioprinting platform incorporated with a functional hydrogel system to study dynamic metastatic cell behaviors *in vitro* (Utama et al., 2020). This 3D bioprinting platform presented herein exhibits several notable advantages over existing approaches for use in versatile biological applications; (1) The 3D bioprinting platform automates multi-well plate printing, thereby facilitating the generation of 3D cancer cell models in a simple, reproducible, viable and HTP manner. (2) The HTP 3D platform allows for the visualization of metastasis markers *in situ* and also for the simultaneous examination of cell movement and the effects of inhibitors. (3) The fine tunability of the hydrogel systems used in the bioprinting platform helps identify cell type specific matrix conditions. (4) Finally, the design of bioprinted hydrogels with sites for proteolytic breakdown enables cells to be retrieved from the hydrogels for downstream molecular analysis post-bioprinting.

Here we utilized the 3D bioprinting platform that combines a fly-by bioprinting technology and a tunable hydrogel system. This approach enables the instant encapsulation and generation of multiple 3D models of different cells and/or matrix components in a well plate for various biological applications. A study has recently developed a HTP bioprinting platform using GelMA hydrogels and a digital light processing-based system (Hwang et al., 2021). Similar to the bioprinting platform presented in this study, their platform was capable of printing for HTP *in situ* fabrication of up to 96 samples per batch. While application of the platform was demonstrated to be a drug-response assay in their study, we further advanced the biological applicability of the HTP bioprinted platform beyond drug screening, including *in situ* and detailed molecular analysis and real-time cell migration tracking. To mimic the ECM, we functionalized the PEG-4MAL hydrogels with peptides containing a fibronectin derived ligand (RGD) for cell adhesion (Singh et al., 2014, Pierschbacher and Ruoslahti, 1984). The addition of MMP-sensitive sites in the hydrogel systems enabled cells to migrate and invade through the hydrogels. The *in situ* analysis demonstrated that the MMP2 and MMP9 proteases were localized on the surface of invasive cancer cells, which may facilitate their proteolytic activation that in turn affects various cell functions, such as proliferation and migration (Brooks et al., 1996, Murphy and Nagase, 2011).

We selected the stiffness values of 0.7 or 1.1 kPa rheology storage modulus values, equivalent to Young’s modulus values (E) of approximately 1.5 or 3.2 kPa respectively as calculated using the formula previously reported (Polio et al., 2018). These were selected as they are close to the stiffness levels of malignant breast tumors (0.2-2.5 kPa (E) (Acerbi et al., 2015, Levental et al., 2009) and ovarian cancers (~3 ± 2.5 kPa (E) (Hopkins et al., 2021)), indicating that the hydrogels were tuned in a biomimetic way. Interestingly, while for the invasive cancer cell lines the effect of the matrix stiffness on cell growth and morphology was trivial, we observed that the migration of those cells was promoted in the stiffer bioprinted matrix. These data are in line with previous studies which reported that increasing matrix stiffness has been shown to induce malignant phenotypes (Acerbi et al., 2015, Kai et al., 2016). Cancer cells can sense matrix stiffening through integrins that regulate cell migration via the activation of various signaling pathways, including the Rho/ROCK pathway (Costa et al., 2013, Kai et al., 2016). Indeed, the inhibitory effect of the ROCK inhibitor on MDA-MB-231 and H1299 cell migration within the 3D matrix could be monitored and quantitated using our 3D bioprinting platform, suggesting its potential application for pre-clinical drug screening for metastatic diseases. Further by taking advantage of the use of proteolytic degradable hydrogel systems in this platform, the downstream molecular analysis using retrieved cells can be performed to better understand the mechanisms of actions for novel drugs or inhibitors.

In summary, here we report multiple capabilities of a HTP 3D bioprinting platform coupled with a highly tunable hydrogel system. This study describes not only the bioprinting technology that can engineer the 3D tumor models mimicking the dynamic tumor microenvironment in a HTP manner, but also the versatile and flexible applications of the bioprinting platform in cancer research. Therefore, we believe that the platform can provide a powerful tool for the study of cell migration and invasion in a 3D environment that mimics tumor growth.

### Significance

3D bioprinting provides a unique approach to the fabrication of complex tissue constructs *in vitro*, yet the use of 3D bioprinted cell models for biological applications has not been fully explored. Here we present a versatile HTP 3D bioprinting approach to study cell movement within the ECM mimiking hydrogel system that enables the biological characterisation of cells. This approach of combining the rapid HTP bioprinting platform and its biological applications has the significant potential for better understanding cell migration and invasion processes and the identification of novel anti-metastasis drugs.

## Acknowledgments

This work was supported by the Children’s Cancer Institute, which is affiliated with the University of New South Wales (UNSW Sydney), and the Sydney Children’s Hospital Network and by grants from the Australian Research Council (ARC) Linkage Grant (LP170100623 to J.J.G., M.K., and J.C.C.R.); the National Health and Medical Research Council (NHMRC) Program Grant (APP1091261 to M.K. and J.J.G.); the NHMRC Principal Research Fellowship (APP1119152 to M.K.), the NHMRC Investigator (APP1196648 to J.J.G.), and grant support from the ARC Centre of Excellence in Convergent Bio-Nano Science and Technology (CE140100036 to J.J.G. and M.K.). The authors would like to thank the Katharina Gaus Light Microscopy, at the Mark Wainwright Analytical Center at UNSW Sydney for their support and resources involved in this work

## Author contributions

Conceptualisation: M.K., J.J.G., and M.J.; Methodology: M.K., J.J.G., M.J., M.E., R.H.U., J.C.C.R., A.P.O.M., A.S, and A.V.; Software: A.S. and A.P.O.M.; Investigation: M.J., J.N.S., M.A.K.T., E.D.; Writing – Original Draft: M.J.; Writing – Review & Editing: M.K., J.J.G., and M.J.; Visualisation: M.J. and J.N.S.; Resources: J.C.C.R., M.E., R.H.U., A.V., A.P.O.M. and A.S.; Funding Acquisition: M.K., J.J.G. and J.C.C.R.

## STAR methods

### Cell lines

MCF7 and H1299 cells were purchased from ATCC. MDA-MB-231 cells were kindly provided by Dr. Rose Boutros (Kids Research, Sydney, NSW, Australia). HEY was a generous gift from Georgia Chenevix-Trench (QIMR Berghofer, Brisbane, Australia). All cells were cultured in either DMEM (MCF7 and MDA-MB-231) or RPMI1640 (HEY and H1299) media supplemented with 10 % FCS. Cells were maintained in a humidified atmosphere containing 5 % CO2 at 37 °C and were mycoplasma free. All cell lines were authenticated using short tandem repeat profiling at Cell Bank Australia and Kinghorn center for clinical genomics, Australia within the last two years.

### Bioprinting of 3D cell laden models

Bioinks and activators for 4 hydrogel combinations were purchased from Inventia Life Science, Sydney, Australia (Cat No. Px01.00, Px01.03P, Px02.00 and Px02.03P for 0.7 kPa, 0.7 kPa+RGD, 1.1 kPa and 1.1 kPa+RGD hydrogels respectively). 3D cell models were printed using the Rastrum 3D bioprinter (Inventia Life Science) as previously described (Utama et al., 2021). Briefly, the structure design and printing protocol were first created using RASTRUM Cloud (Inventia Life Science). Cells were primed (1 × 10^6^ cells for MCF7, MDA-MB-231 and H1299 with a final seeding density of ~ 500 cells per well and 0.5 × 10^6^ cells for HEY with a final seeding density of ~ 250 cells per well) prior to being printed within Small Plug or Imaging Model. Cells were also bioprinted using Large Plug for molecular downstream analysis (qPCR and Western blotting) available through RASTRUM Cloud. All printing was conducted in flat bottom 96-well plates (Corning). In parallel, Matrigel-encapsulated cells were manually prepared in a 96 well plate as a control. 500 cells (MCF7, MDA-MB-231 and H1299) or 250 cells (HEY) were mixed in 2 μl of growth-factor reduced Matrigel solution (In Vitro Technologies), which was 1:1 diluted with cell culture media.

### Live and dead cell staining

Cell viability analysis was performed using Live/Dead viability/cytotoxicity kit, for mammalian cells (Invitrogen, Cat No. L3224) according to the manufacturer’s instructions. Briefly, cells were bioprinted in multi-well plates and cultured for 7 days post-printing. At day 7, cells were rinsed with DPBS and stained with 100 μL of live/dead stock solutions (10 μM Ethidium Homodimer-1 (EthD1) and 5 μM Calcein AM in DPBS) and incubated for 30 min. Images were taken at 5X magnification using green fluorescence channel for live cells and red fluorescence channel for dead cells using Celldiscoverer 7 (Zeiss). Images were analysed and visualized uisng Arivis 4D software.

### Alamar Blue cell proliferation assay

Cells were bioprinted with each hydrogel condition in a 96 well plate and cultured for up to 7 days. Cells were incubated with resazurin-based reagent (Sigma-Aldrich) at 10 % of media volume, for 16 h at time-points of day 1, 2, 3 and 7. The assay was read with Benchmark Plus plate reader (BIO-RAD) at 570-595 nm and percent viability normalized to ethanol treated cells (negative control).

### Immunofluorescent staining

Cells were bioprinted in a 96 well plate and cultured for up to 7 days. At day 7, cells were fixed and permeabilized in 4 % Paraformaldehyde/0.1 % Triton-X100/DPBS solution in the plate for 2 h at room temperature followed by blocking with 5 % BSA/TBST overnight at 4 °C. After removing the blocking solution, cells were incubated with primary antibodies including anti-E-Cadherin (Cat No. 14472, Cell Signaling Technology, 1:500 dilution), anti-Vimentin (Cat No. 5741, Cell Signaling Technology, 1:500 dilution), anti-MMP2 (Cat No. ab37150, Abcam, 1:200 dilution) and anti-MMP9 (Cat No. ab38898, Abcam, 1:200 dilution) overnight at 4 °C. All antibodies were diluted with 5 % BSA/TBST. Each well was rinsed twice with TBST then cells were incubated with Alexa Fluor 488 labelled secondary antibodies (anti-rabbit IgG (Cat No. A11034, Invitrogen, 1:500 dilution) or anti-mouse IgG (Cat No. A32723, Invitrogen, 1:500 dilution)) overnight at 4 °C. For F-actin staining, bioprinted cells were incubated with phalloidin conjugated to Alexa Fluor^®^ 568 (Cat No. A12380, Life Technologies, 1:200 dilution) on a shaker 2 h at room temperature. After washing with TBST twice, nuclei were counterstained with 0.2 mM Hoechst 33342 (Cat No. 62249, ThermoFisher Scientific, stock concentration 20 mM) in DPBS for 10 minutes at room temperature. The 3D bioprinted cells were then imaged using the Leica TCS SP8 DLS (Digital LightSheet) confocal microscope. Images were taken using 10X objective, Argon laser (458 and 488nm) and DPSS 561 Laser (561 nm) (Inset zoomed-in images: 2X or 4X zoom). Z-stacks were defined from the bottom to the top of the 3D hydrogels and an image stack on the z-axis was taken per well.

### Cell recovery from 3D bioprinted matrices

RASTRUM Cell Retrieval Solution was purchased from Inventia Life Sciences (Cat No.F235). Cells were bioprinted with each hydrogel condition in a 96 well plate and cultured for 7 days. Cells were retrieved and pellets were collected according to manufacturer’s instructions. Briefly, cells were rinsed with DPBS and incubated with 75 μl Cell Retrieval Solution for 20 min at 37 °C/5 % CO_2_. Solution was collected after pipetting up and down in each well several times, then wells were washed with DPBS and contents were combined with collected cell suspension. Cells were pelleted at 1200 rpm and pellets rinsed twice with cold DPBS. Final cell pellets were snap frozen on dry ice and stored at −80 °C for further analysis.

### Real-time qPCR (qPCR)

Cells were bioprinted with each hydrogel condition in a 96 well plate and cultured for 7 days before cells were extracted using method described above. RNA was extracted from cell pellets using QIAGEN RNeasy Mini Kit (Cat No. 74104) according to manufacturer’s instructions. RNA concentration was determined using ThermoScientific Nanodrop 2000 Spectrophotemeter quantification. RNA was reverse transcribed using MMLV reverse transcriptase (Life Technologies) and qPCR was performed with the 7900HT Fast Real-Time PCR System (Life Technologies) as previously described (Jung et al., 2017). Briefly, qPCR was set up using 2 μg cDNA and KAPA Probe Fast master mix (Roche, Cat No. KK4705). For target genes, TaqMan^™^ Gene Expression Assay probes for CDH1 (Hs01023895_m1), VIM (Hs00185584_m1), MMP2 (Hs01548727_m1) and MMP9 (Hs00957562_m1) were used to measure gene expression levels and normalized against GAPDH housekeeping gene (Hs02786624_g1). For each target gene, expression level was quantified in relation to the expression of a control gene using the ΔΔCt method to provide relative quantification. Gene expression values normalized to each control gene were calculated as the average of the expression value for each target gene.

### Western blot analysis

Western blot was performed as previous described (Gao et al., 2020). Briefly, lysates were made from cell pellets using RIPA buffer containing 1 mM EDTA and 10 % Protease Inhibitor cocktail (Sigma-Aldrich). Protein concentration was measured using Pierce^™^ BCA Protein assay (ThermoFisher Scientific, Cat No. 23225) as per manufacturer’s instructions. 20 μg protein was run on 4-15 % Mini-PROTEAN TGX Stain-Free Protein Gels (Bio-Rad, Cat No. 4568085) before being transferred to nitrocellulose membranes. Antibodies include anti-E-cadherin (Cat No. 14472, Cell Signaling Technology, 1:1000 dilution) and anti-Vimentin (Cat No. 5741, Cell Signaling Technology, 1:1000 dilution), GAPDH (Cat No. ab8245, Abcam, 1:100,000 dilution) primary antibodies and anti-rabbit-HRP (Cat No. P0448, DAKO, 1:5000 dilution) and anti-mouse (Cat No. P0447. DAKO, 1:5000 dilution) secondary antibodies. All antibodies were diluted in 5 % skim milk/TBST. Signals were detected using Clarity ECL Western Blotting Substrate (BIO-RAD).

### Live cell tracking

Cells were bioprinted with each hydrogel condition in a glass-bottom 96 well plate. Some wells were treated with 80 % ethanol for cell death and used as a control to correct for non-cell movement. The distance and speed values in live cells below 75 % of those in the control wells were considered to be artifacts and excluded in the analysis. The plates were incubated at 5 % CO_2_ and 37 °C for 3 h prior to adding a ROCK inhibitor, Y-27632 (Cat No. 72302, StemCell Technologies) and global myosin inhibitor, blebbistatin (Cat No. B0560, Sigma-Aldrich) were made up to 10 mM in DMSO. Bioprinted drugged plates were incubated for 24 h at 37 °C/5 % CO_2_. Incucyte^®^ Nuclight Rapid Red Dye (1:1000 in culture media, Sartorius, Cat No. 4717) was added with fresh drugged media 1 h prior to cell tracking. Z-stack images were taken every 15 min for 24 h on Zeiss CellDiscoverer 7 microscope using bright field and red fluorescence channel at 10X objective. The microscope incubator was set to 37 °C and 5 % CO_2_. A minimum of 95 cell tracks per well were included in the analysis. Cell tracking analyses were quantified using a built-in tracking algorithm in Imaris software (Bitplane, Zurich, Switzerland).

### Statistical analysis

Statistical analyses were performed using the GraphPad Prism v9.0 software (GraphPad Software). For the cell movement analysis, unpaired, two-tailed Mann-Whitney tests were used to determine statistical differences between two different groups (cell types or gel conditions). For comparison of multiple samples, Kruskal–Wallis one-way analysis with a post hoc Dunn test was used.

## Supplemantal information

**Supplementary Figure 1.**
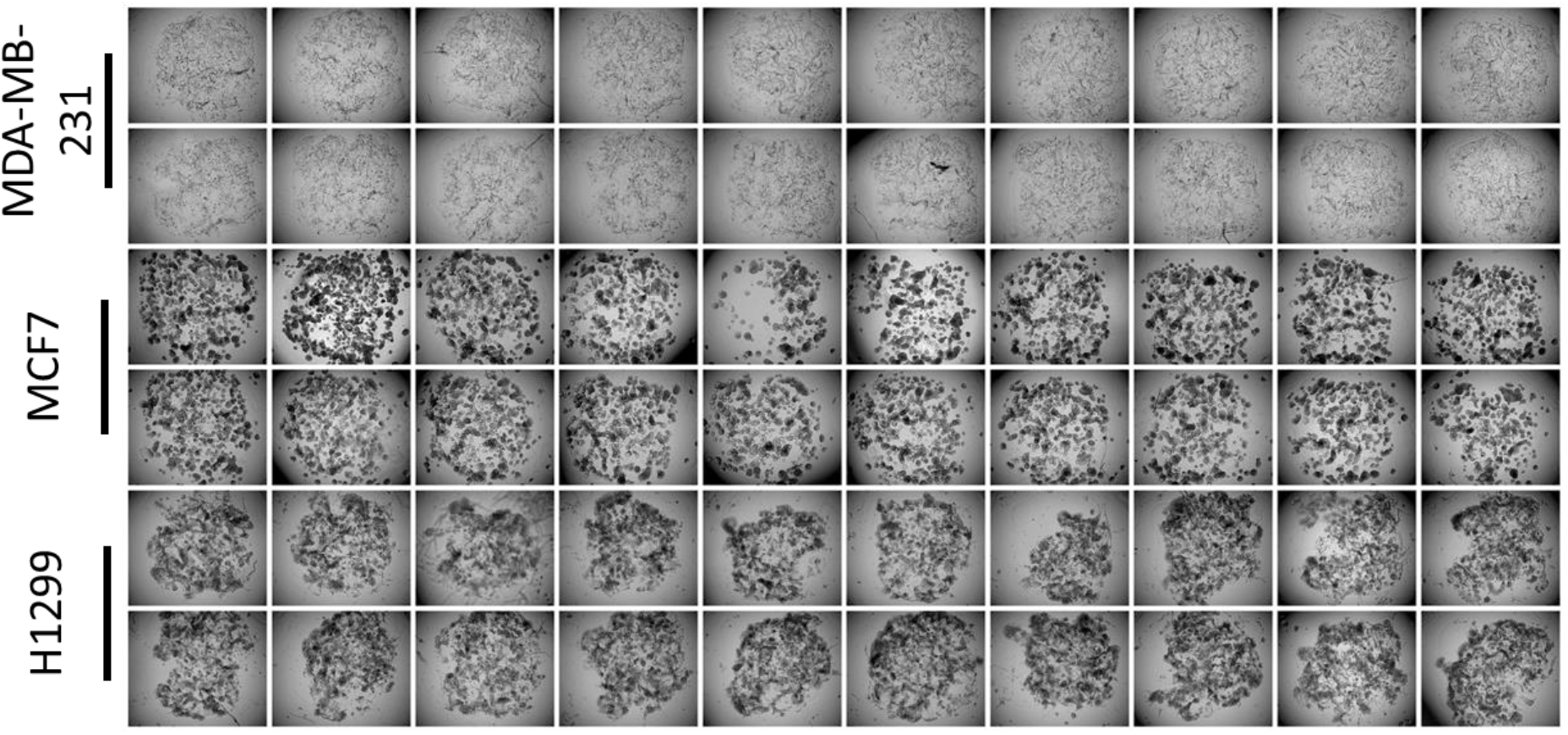
Generation of multiple cell models in a 96 well plate. The HTP 3D bioprinting platform allows to print up to 3 different cancer models in a multi-well plate. MDA-MB-231, MCF7 and H1299 cells were bioprinted with 0.7 kPa+RGD hydrogels in the inner 60 wells of a 96 well plate. The plate was incubated at 37°C for 7 days. Bright-field images were taken at a single plane (5X objective).

**Supplementary Figure 2.**
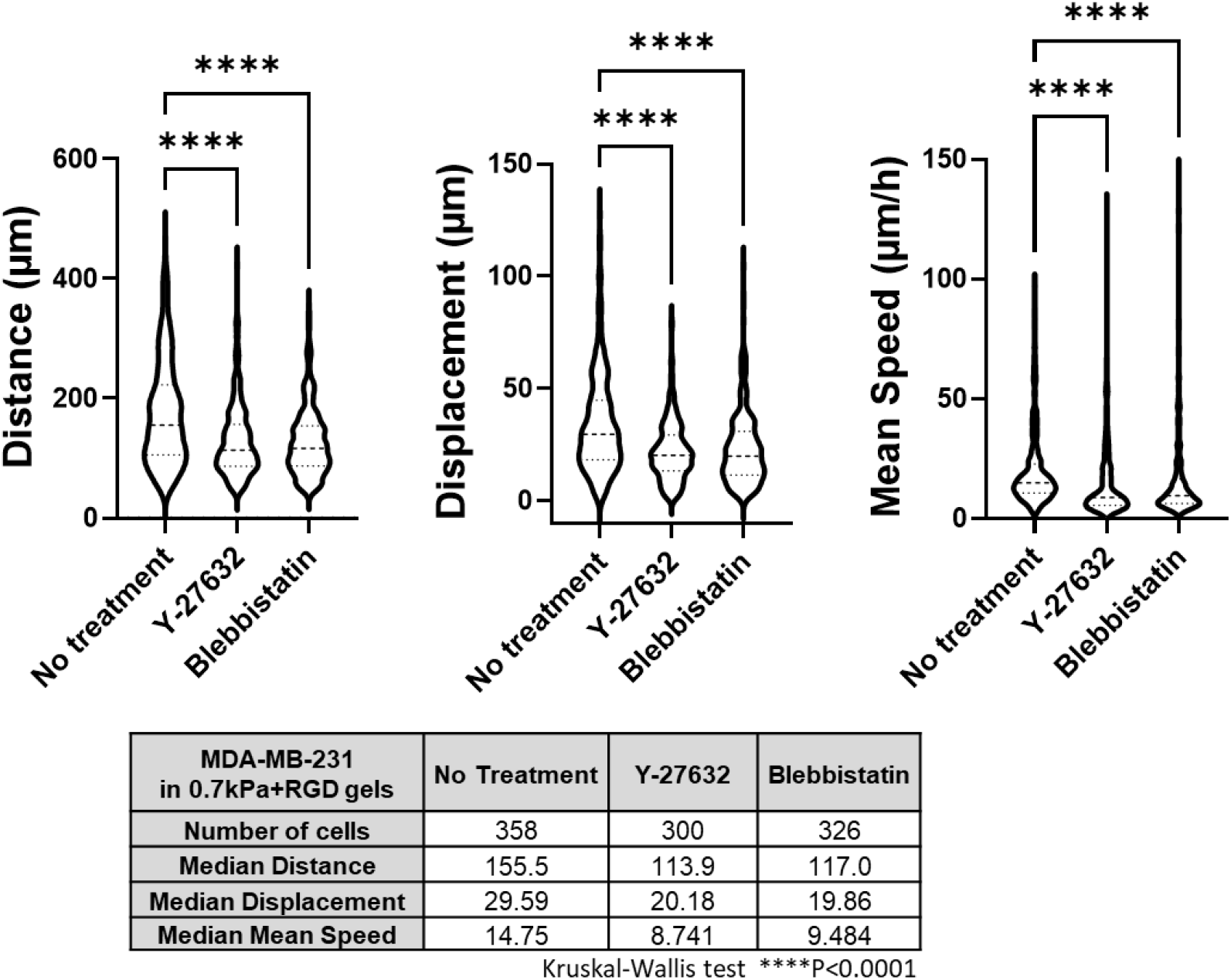
Chemical inhibitors of MDA-MB-231 cell migration within 0.7 kPa+RGD hydrogels. MDA-MB-231 cells were bioprinted in 0.7 kPa+RGD hydrogels. Either Y-27632 (a ROCK inhibitor) or Blebbistatin (a global myosin inhibitor) were treated in different wells in a 96 well plate. 3D cell movement of MDA-MB-231 cells in the absence or presence of the inhibitors was compared and quantitated. Kruskal–Wallis one-way analysis with a post hoc Dunn test was performed. ****P<0.0001

## Supplementary Movies

Distinct cell morphology and movement patterns of two variants of breast cancer cells in 3D matrices. All images were taken with IncuCyte microscopy (objective 4X for Matrigel and 10X for bioprinted).

Movie 1. MCF7 in Matrigel

Movie 2. MDA-MB-231 in Matrigel

Movie 3. MCF7 in a 0.7 kPa bioprinted gel

Movie 4. MDA-MB-231 in a 0.7 kPa+RGD bioprinted gel

## References

Acerbi, I., Cassereau, L., Dean, I., Shi, Q., Au, A., Park, C., Chen, Y. Y., Liphardt, J., Hwang, E. S. & Weaver, V. M. 2015. Human breast cancer invasion and aggression correlates with ECM stiffening and immune cell infiltration. Integrative Biology, 7, 1120–1134.

Amaral, R. L. F., Miranda, M., Marcato, P. D. & Swiech, K. 2017. Comparative Analysis of 3D Bladder Tumor Spheroids Obtained by Forced Floating and Hanging Drop Methods for Drug Screening. Frontiers in physiology, 8, 605–605.

Benton, G., Kleinman, H. K., George, J. & Arnaoutova, I. 2011. Multiple uses of basement membrane-like matrix (BME/Matrigel) in vitro and in vivo with cancer cells. International Journal of Cancer, 128, 1751–1757.

Brabletz, T., Kalluri, R., Nieto, M. A. & Weinberg, R. A. 2018. EMT in cancer. Nature Reviews Cancer, 18, 128–134.

Brooks, P. C., StrÖmblad, S., Sanders, L. C., Von Schalscha, T. L., Aimes, R. T., Stetler-Stevenson, W. G., Quigley, J. P. & Cheresh, D. A. 1996. Localization of Matrix Metalloproteinase MMP-2 to the Surface of Invasive Cells by Interaction with Integrin αvβ3. Cell, 85, 683–693.

Costa, P., Scales, T. M. E., Ivaska, J. & Parsons, M. 2013. Integrin-Specific Control of Focal Adhesion Kinase and RhoA Regulates Membrane Protrusion and Invasion. PLOS ONE, 8, e74659.

Ding, H., Illsley, N. P. & Chang, R. C. 2019. 3D Bioprinted GelMA Based Models for the Study of Trophoblast Cell Invasion. Scientific Reports, 9, 18854.

Fang, Y. & Eglen, R. M. 2017. Three-Dimensional Cell Cultures in Drug Discovery and Development. SLAS discovery: advancing life sciences R & D, 22, 456–472.

Friedl, P. & Wolf, K. 2003. Tumour-cell invasion and migration: diversity and escape mechanisms. Nature Reviews Cancer, 3, 362–374.

Gao, J., Jung, M., Mayoh, C., Venkat, P., Hannan, K. M., Fletcher, J. I., Kamili, A., Gifford, A. J., Kusnadi, E. P., Pearson, R. B., et al. 2020. Suppression of ABCE1-Mediated mRNA Translation Limits N-MYC-Driven Cancer Progression. Cancer Research, 80, 3706.

Havel, L. S., Kline, E. R., Salgueiro, A. M. & Marcus, A. I. 2015. Vimentin regulates lung cancer cell adhesion through a VAV2-Rac1 pathway to control focal adhesion kinase activity. Oncogene, 34, 1979–1990.

Heinrich, M. A., Bansal, R., Lammers, T., Zhang, Y. S., Michel Schiffelers, R. & Prakash, J. 2019. 3D-Bioprinted Mini-Brain: A Glioblastoma Model to Study Cellular Interactions and Therapeutics. Advanced Materials, 31, 1806590.

Hopkins, T. I. R., Bemmer, V. L., Franks, S., Dunlop, C., Hardy, K. & Dunlop, I. E. 2021. Mapping the mechanical microenvironment in the ovary. bioRxiv, 2021.01.03.425098.

Hwang, H. H., You, S., Ma, X., Kwe, L., Victorine, G., Lawrence, N., Wan, X., Shen, H., Zhu, W. & Chen, S. 2021. High throughput direct 3D bioprinting in multiwell plates. Biofabrication, 13, 025007.

Jung, M., Russell, A. J., Liu, B., George, J., Liu, P. Y., Liu, T., Defazio, A., Bowtell, D. D. L., Oberthuer, A., London, W. B., et al. 2017. A Myc Activity Signature Predicts Poor Clinical Outcomes in Myc-Associated Cancers. Cancer Research, 77, 971.

Kai, F., Laklai, H. & Weaver, V. M. 2016. Force Matters: Biomechanical Regulation of Cell Invasion and Migration in Disease. Trends in Cell Biology, 26, 486–497.

Kratochvil, M. J., Seymour, A. J., Li, T. L., PaŞca, S. P., Kuo, C. J. & Heilshorn, S. C. 2019. Engineered materials for organoid systems. Nature Reviews Materials, 4, 606–622.

Levental, K. R., Yu, H., Kass, L., Lakins, J. N., Egeblad, M., Erler, J. T., Fong, S. F. T., Csiszar, K., Giaccia, A., Weninger, W., et al. 2009. Matrix Crosslinking Forces Tumor Progression by Enhancing Integrin Signaling. Cell, 139, 891–906.

Liu, C.-Y., Lin, H.-H., Tang, M.-J. & Wang, Y.-K. 2015. Vimentin contributes to epithelial-mesenchymal transition cancer cell mechanics by mediating cytoskeletal organization and focal adhesion maturation. Oncotarget, 6, 15966–15983.

Lutolf, M. P., Lauer-Fields, J. L., Schmoekel, H. G., Metters, A. T., Weber, F. E., Fields, G. B. & Hubbell, J. A. 2003. Synthetic matrix metalloproteinase-sensitive hydrogels for the conduction of tissue regeneration: engineering cell-invasion characteristics. Proceedings of the National Academy of Sciences of the United States of America, 100, 5413–5418.

Mehra, A., Lee, K. H. & Hatzimanikatis, V. 2003. Insights into the relation between mRNA and protein expression patterns: I. theoretical considerations. Biotechnology and Bioengineering, 84, 822–833.

Meng, F., Meyer, C. M., Joung, D., Vallera, D. A., Mcalpine, M. C. & Panoskaltsis-Mortari, A. 2019. 3D Bioprinted In Vitro Metastatic Models via Reconstruction of Tumor Microenvironments. Advanced Materials, 31, 1806899.

Moroni, L., Burdick, J. A., Highley, C., Lee, S. J., Morimoto, Y., Takeuchi, S. & Yoo, J. J. 2018. Biofabrication strategies for 3D in vitro models and regenerative medicine. Nature reviews. Materials, 3, 21–37.

Murphy, G. & Nagase, H. 2011. Localizing matrix metalloproteinase activities in the pericellular environment. The FEBS journal, 278, 2–15.

Noshadi, I., Hong, S., Sullivan, K. E., Shirzaei Sani, E., Portillo-Lara, R., Tamayol, A., Shin, S. R., Gao, A. E., Stoppel, W. L., Black Iii, L. D., et al. 2017. In vitro and in vivo analysis of visible light crosslinkable gelatin methacryloyl (GelMA) hydrogels. Biomaterials Science, 5, 2093–2105.

Novikov, N. M., Zolotaryova, S. Y., Gautreau, A. M. & Denisov, E. V. 2021. Mutational drivers of cancer cell migration and invasion. British Journal of Cancer, 124, 102114.

Pierschbacher, M. D. & Ruoslahti, E. 1984. Cell attachment activity of fibronectin can be duplicated by small synthetic fragments of the molecule. Nature, 309, 30–33.

Polio, S. R., Kundu, A. N., Dougan, C. E., Birch, N. P., Aurian-Blajeni, D. E., Schiffman, J. D., Crosby, A. J. & Peyton, S. R. 2018. Cross-platform mechanical characterization of lung tissue. PLOS ONE, 13, e0204765.

Roche, J. 2018. The Epithelial-to-Mesenchymal Transition in Cancer. Cancers, 10, 52.

Singh, S. P., Schwartz, M. P., Lee, J. Y., Fairbanks, B. D. & Anseth, K. S. 2014. A peptide functionalized poly(ethylene glycol) (PEG) hydrogel for investigating the influence of biochemical and biophysical matrix properties on tumor cell migration. Biomaterials Science, 2, 1024–1034.

Swaminathan, S., Hamid, Q., Sun, W. & Clyne, A. M. 2019. Bioprinting of 3D breast epithelial spheroids for human cancer models. Biofabrication, 11, 025003.

Swaminathan, V., Mythreye, K., Brien, E. T., Berchuck, A., Blobe, G. C. & Superfine, R. 2011. Mechanical Stiffness Grades Metastatic Potential in Patient Tumor Cells and in Cancer Cell Lines. Cancer Research, 71, 5075.

Utama, R. H., Atapattu, L., O’mahony, A. P., Fife, C. M., Baek, J., Allard, T., O’mahony, K. J., Ribeiro, J. C. C., Gaus, K., Kavallaris, M., et al. 2020. A 3D Bioprinter Specifically Designed for the High-Throughput Production of Matrix-Embedded Multicellular Spheroids. iScience, 23, 101621.

Utama, R. H., Tan, V. T. G., Tjandra, K. C., Sexton, A., Nguyen, D. H. T., O’Mahony, A. P., Du, E. Y., Tian, P., Ribeiro, J. C. C., Kavallaris, M., et al. 2021. A Covalently Crosslinked Ink for Multimaterials Drop-on-Demand 3D Bioprinting of 3D Cell Cultures. Macromolecular Bioscience, n/a, 2100125.

Weigelt, B. & Bissell, M. J. 2008. Unraveling the microenvironmental influences on the normal mammary gland and breast cancer. Seminars in cancer biology, 18, 311–321.

Zhou, X., Zhu, W., Nowicki, M., Miao, S., Cui, H., Holmes, B., Glazer, R. I. & Zhang, L. G. 2016. 3D Bioprinting a Cell-Laden Bone Matrix for Breast Cancer Metastasis Study. ACS Applied Materials & Interfaces, 8, 30017–30026.

